# Effects of green seaweed extract on *Arabidopsis* early development suggest roles for hormone signalling in plant responses to algal fertilisers

**DOI:** 10.1101/400143

**Authors:** Fatemeh Ghaderiardakani, Ellen Collas, Deborah Kohn Damiano, Katherine Tagg, Neil S. Graham, Juliet C. Coates

**Author notes:** Equal author contribution (Ellen Collas, Deborah Kohn Damiano, Katherine Tagg).

## Abstract

The growing population requires sustainable, environmentally-friendly crops. The plant growth-enhancing properties of algal extracts have suggested their use as biofertilisers. The mechanism(s) by which algal extracts affect plant growth are unknown.

We examined the effects of extracts from the common green seaweed *Ulva intestinalis* on germination and root development in the model land plant *Arabidopsis thaliana*. *Ulva* extract concentrations above 0.1% inhibited *Arabidopsis* germination and root growth. *Ulva* extract <0.1% stimulated root growth. All concentrations of *Ulva* extract inhibited lateral root formation. An abscisic-acid-insensitive mutant, *abi1*, showed altered sensitivity to germination- and root growth-inhibition inhibition. Ethylene- and cytokinin-insensitive mutants were partly insensitive to germination-inhibition. This suggests that different mechanisms mediate each effect of *Ulva* extract on early *Arabidopsis* development and that multiple hormones contribute to germination-inhibition.

Elemental analysis showed that *Ulva* contains high levels of Aluminium ions (Al^3+^). Ethylene and cytokinin have been suggested to function in Al^3+^-mediated root growth inhibition: our data suggest that if *Ulva* Al^3+^ levels inhibit root growth, this is via a novel mechanism. We suggest algal extracts should be used cautiously as fertilisers, as the inhibitory effects on early development may outweigh any benefits if the concentration of extract is too high.

## Introduction

Plant growth, development and productivity is affected by various abiotic (physical) and biotic (biological) factors. Responses to these factors determine cropping pattern and plant distribution ^1^. Global demand for crops is predicted to increase ∼100% from 2005 to 2050, while ∼795 million people worldwide were undernourished in 2014–16 ^2,3^.

Current global food challenges and pressure on the food production industry are due to the exponentially growing human population and increasing soil- and water issues compounding the pressure induced by anthropogenic climate change. The frequency of abiotic environmental stresses (flooding, drought, water limitation, salinity and extreme temperatures) is increasing ^4^ and causing crop losses worldwide ^5-7^. More intense, frequent droughts in Africa, southern South America and southern Europe and increased flooding in temperate regions will drive future crop yield decline ^8-10^. Soil salinity threatens agriculture and natural ecosystems ^11-13^. Intensive farming leads to unfavourable conditions for crop growth, development and survival ^14^.

Humans have used seaweeds (macroalgae) and seaweed-based products for centuries, for food, fuel, aquaculture, cosmetics, colouring dyes and therapeutic/botanical applications ^14-16^. The earliest written reference to using seaweed as a fertiliser is from Roman times ^17^. Applying seaweeds/seaweed extracts in modern agriculture leads to increased seed germination rates, improved plant development (flowering, leaf quality and root system architecture), elevated defence against pathogens and pests ^18^ and protection against nutrient deficiency and environmental stresses including salinity ^19^, cold or drought ^20-23^. Seaweed fertilisers have been used in agricultural programs to improve soil management, disease management, nutritional strategies, water efficiency and drought tolerance ^23^.

Several manufacturing practices are used to liquefy seaweed biomass ^21,23,24^. Seaweed extracts are marketed as liquid biofertilisers or biostimulants containing a variety of plant growth-promoting components – those identified include plant growth regulators (phytohormones), minerals and trace elements, quaternary ammonium molecules (e.g. betaines and proline), polyuronides (e.g. alginates/fucoidans) and lipid-based molecules e.g. sterols ^23^. Seaweed products are also available in soluble powder form. Depending on whether algal extract is applied as liquid fertiliser or seaweed manure to plant roots, or as a leaf spray, different plant responses to seaweeds occur ^14,21^. The mechanism by which seaweed fertilisers affect plant growth, development and yield is currently unknown. Crop plants treated with seaweed extracts showed similar physiological responses to those treated with plant growth-regulatory substances ^20^. Phytohormones detected in seaweed extracts are auxins, cytokinins, gibberellins, abscisic acid and brassinosteroids ^25-27^ but chemical components other than phytohormones, which elicit physiological responses reminiscent of plant hormones, have also been detected ^28^. One hypothesis is that the effects of seaweed fertilisers are due either directly or indirectly to phytohormones: seaweed extracts may themselves contain beneficial phytohormones, or may contain substances that trigger land plant signaling pathways that usually respond to these signals. Which, if either, of these scenarios occurs is not clear.

Although seaweeds could potentially benefit plant growth by providing macronutrients, including nitrogen (N), phosphorus (P), ammonium (NH4+) and potassium (K), studies have consistently shown that seaweed extracts’ beneficial effects are not due to macronutrients, particularly at the concentrations used in the field ^20,29^. Very dilute seaweed extracts (1:1000 or below) still have biological activity but the compound(s) involved are unknown: the beneficial effects may involve several plant growth-promoters working synergistically ^25,30-32^.

Understanding at a mechanistic level how seaweed fertilisers affect land plant growth and development is important. Previous studies have applied a diverse range of extracts from brown, green and red seaweeds to a heterogeneous range of crop plants ^33,34^. Generally, lower concentrations of an algal extract have beneficial effects on root- and shoot growth while higher concentrations have inhibitory effects ^34-38^. Thus, algal extract concentration is critical to its effectiveness. However, because of the range of plants, seaweeds and extraction methods used, “positive” concentrations of algal extract ranged from 0.002%-0.2% while inhibitory concentrations ranged from 0.1%-1%.

In this paper, we establish a “standardised” laboratory-based system to determine the molecular mechanisms by which seaweeds can affect land plant productivity, using model organisms. The extensively-studied model plant *Arabidopsis* ^39^, from *Brassicaceae* (cabbage) family, was the first plant with a sequenced genome ^40^ and extensive mutant collections are available, including mutants in hormone signaling and perception ^41-45^. Phytohormone biosynthetic/signalling pathways have been determined, yielding a broad understanding of plant responses to stimuli ^46-48^. Employing *Arabidopsis* as a model organism has enabled translation of the understanding of plant growth and development to crops and agriculture ^49-51^.

The green seaweed *Ulva* (sea lettuce; green nori) is an emerging experimentally-tractable model organism to study macroalgal development, growth, morphogenesis (reviewed in Wichard 2015). *Ulva* is a cosmopolitan macroalgal genus, the main multicellular branch of the Chlorophyte algae, and the most abundant Ulvophyceae representative ^52,53^. Ulvophyceae are multicellular algae with simple morphology compared to land plants. Distinctive features that make *Ulva* attractive as model systems are the small genome [100-300Mb; ^54,55^] (the established model system *Ulva mutabilis* is currently being sequenced), symbiotic growth with bacterial epiphytes, naturally-occurring developmental mutants (in *Ulva mutabilis*), simple organization of the thallus (body) consisting of three differentiated cell types (blade, stem and rhizoid), laboratory cultivation ^56,57^ and the ability to generate stable transgenic lines ^58-60^.

The species of *Ulva* chosen for this study was *Ulva intestinalis*, an intertidal alga found worldwide, which can be lab-grown similarly to *Ulva mutabilis* ^56,61^. We compared directly the growth- or inhibition parameters of different concentrations of *Ulva intestinalis* extract versus a control, applied to both wild-type and mutant *Arabidopsis* genotypes. By using two experimentally tractable organisms we have begun to understand the plant signalling pathways that can be triggered by algal extract. As *Ulva* genetic manipulation becomes better-established ^59^ this raises the possibility of future engineering of improved macroalgal fertiliser properties.

## Results

### Concentrations of *Ulva* extract of 0.5% and above inhibit wild-type *Arabidopsis* seed germination

Previous experiments demonstrated conflicting effects of different concentrations of algal extract on seed germination, e.g. in tomato ^33,34^. To investigate the effect of *Ulva intestinalis* extract on *Arabidopsis* germination, concentrations of *Ulva* extract ranging from 0 – 1.0% was tested (Fig. 1). All *Ulva* extract concentrations from 0.5% upwards delayed wild-type germination. The final germination percentage was reduced in 0.8% and 1.0% *Ulva* extract: only about half the seeds germinated in 1.0% *Ulva* extract after a week (Fig.1a). Concentrations of 0.3% *Ulva* extract and below had no effect on seed germination and no stimulatory effect of *Ulva* extract on germination was observed at any concentration tested (Fig. 1a).

**Figure 1.**
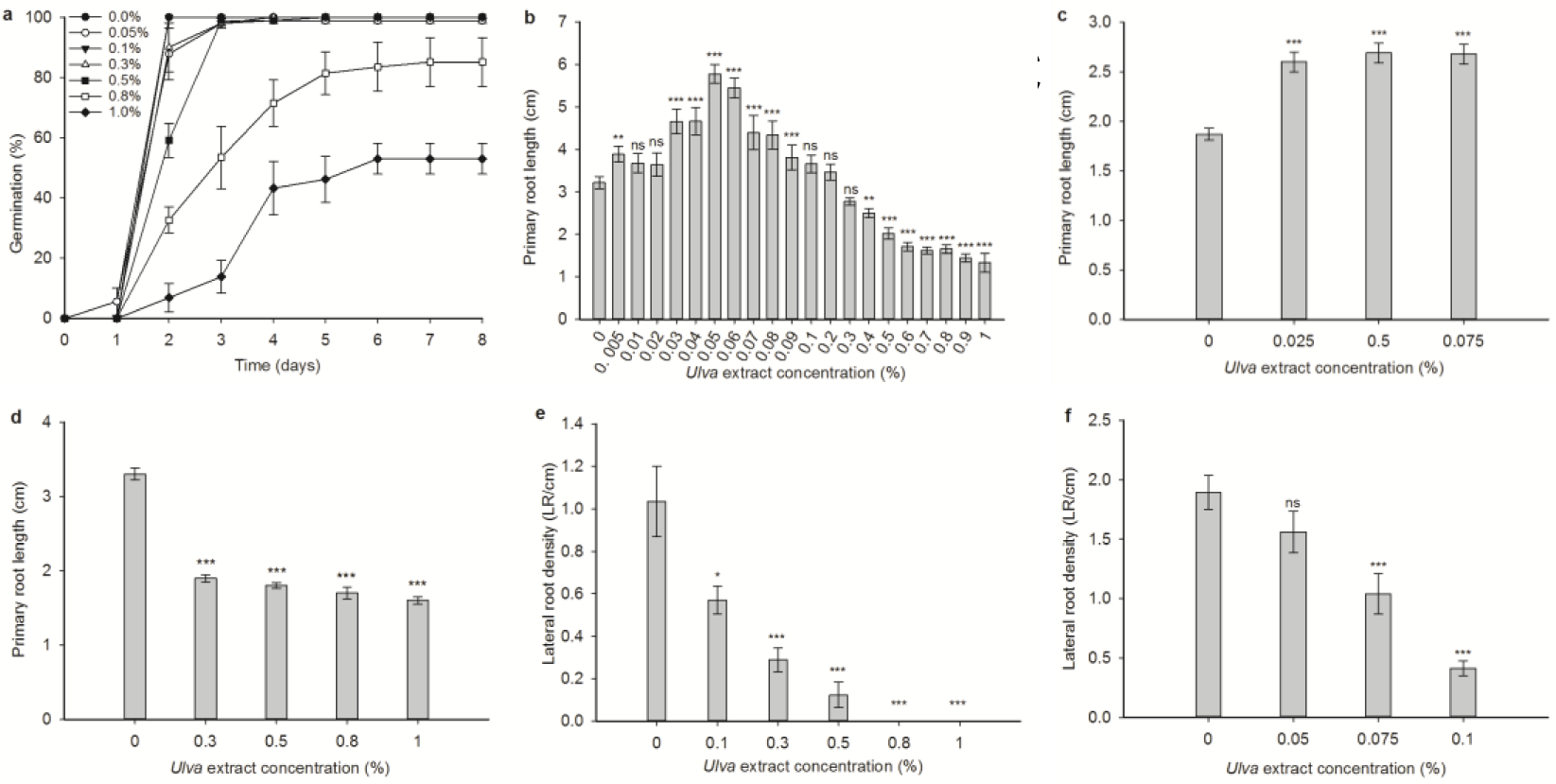
*Ulva* extract inhibits germination and root growth at high concentrations, promotes root growth at lower concentrations, and inhibits lateral root formation, even at concentrations that stimulate primary root growth. (a) Effect of 0-1% *Ulva* extract on wild-type *Arabidopsis* seed germination. Day 3: 0.8% *Ulva* extract significantly (p<0.05% - Kruskal Wallis test) different to control and 0.3% (each p=0.038) and 1% significantly different from control, 0.1% (each p=0008), 0.05% (p=0.025), 0.3% (p=0.035) and 0.5% (p=0.030). Day 4: 0.8% significantly different from control, 0.3% and 0.5% (all p=0.035) and 1% significantly different from control, 0.1% and 0.3% (all p=0.015) and 0.05% (p=0.048). Days 5, 6, 7: control, 0.1%, 0.3%, 0.5% treatments significantly different from 1% (p=0.016 on days 5 and 6; p=0.01 on day 7). (b) Effect of 0-1% *Ulva* extract on wild-type primary root growth in 14-day old seedlings. Significant (p<0.05) differences between treatment and control (Mann-Whitney U-test) were seen with 0.005% (p=0.006), 0.03% (p=0.0001), 0.04% (p=0.0005), 0.05% (p=0.0000), 0.06% (p=0.0000), 0.07% (p=0.0345), 0.08% (p=0.0009), 0.09% (p=0.0000), 0.3% (p=0.0053), 0.4% (p=0.045) and 0.5%-1% (p=0.0000). n=10-40 seedlings per treatment. (c) Primary root growth of wild-type seedlings grown for 10 days on non-nutrient-agar is significantly stimulated by low concentrations (0.025, 0.05 and 0.075%) of *Ulva* extract compared to the control (p=0.0001, p=0.0000, p=0.0000 respectively: t-test). n=30-35 seedlings per treatment. (d) Effect of *Ulva* extract ≥0.3% on wild-type root growth after transferring 3-day old seedlings from control medium, followed by growth for 7 days. Significant inhibition compared to 0% is seen for each concentration (all p=0.0000: t-test). n=30-35 seedlings per treatment. (e) Effect of 0.1-1% *Ulva* extract on lateral root density of wild-type seedlings. Significant inhibition compared to the control (Mann-Whitney U-test) is seen at 0.3% (p=0.0001), 0.5%, 0.8% and 1.0% (all p=0.0000). n=20 seedlings per treatment. (f) Effect of 0.05-0.1% *Ulva* extract on lateral root density of wild-type seedlings. Significant inhibition compared to the 0% control (Mann-Whitney U-test) is seen at 0.75% (p=0.0008) and 0.1% (p=0.0000). n=17-20 seedlings per treatment. All panels: asterisks - significant differences compared to 0% control: * p<0.05, ** p<0.01, *** p<0.001. Bars - standard error of the mean.

### *Ulva* extract stimulates *Arabidopsis* primary root growth at low concentrations and inhibits root growth at higher concentrations

Having demonstrated that seed germination is inhibited by *Ulva* extract, we sought to discover whether the next stage of development, primary root elongation, was also affected by *Ulva* extract. Seeds were germinated, and seedlings grown, on standard growth medium containing a range of *Ulva* extract concentrations ranging from 0 to 2%. *Ulva* extract significantly stimulated root growth at concentrations from 0.03-0.08% (∼80% stimulation at 0.06%), while concentrations of 0.3% and above had an inhibitory effect on root growth (∼68% inhibition at 2%) (Fig. 1b). The stimulatory effect of *Ulva* extract was similarly present when seedlings were grown on non-nutrient-containing agar (Fig. 1c).

To ascertain whether the inhibitory effect of *Ulva* extract concentrations ≥0.3% on root growth was simply a consequence of delayed germination (Fig. 1a), we conducted an experiment where seedlings were germinated on normal growth medium for three days before transferring to medium containing *Ulva* extract. Root growth was once again inhibited by *Ulva* extract, showing that higher concentrations of *Ulva* extract have an inhibitory effect on root growth, independent from any effect on germination (Fig. 1d).

### *Ulva* extract inhibits *Arabidopsis* lateral root formation

Once the *Arabidopsis* primary root is established, it acquires branches, or lateral roots (LRs), as the seedling matures to secure anchorage and extract micro- and macronutrients from the soil ^62,63^. Having ascertained that *Ulva* extract affects primary root growth, we went on to investigate the effect of *Ulva* extract on LR formation. Increasing concentrations of *Ulva* extract show a progressive inhibition in the density of LR branching from the primary root, even at concentrations that stimulate primary root growth (Fig. 1e, f).

In summary, *Ulva* extract inhibits germination, has a biphasic effect on primary root growth (stimulatory at low concentrations; inhibitory at higher concentrations) and inhibits LR formation. Taken together, our data are reminiscent of the effect of the plant hormone abscisic acid (ABA) on germination and early root development, as ABA is a negative regulator of germination ^64^, shows a biphasic effect on primary root growth ^65-67^, and inhibits LR growth at concentrations that stimulate primary root growth but do not affect germination ^68^.

### The germination-inhibitory effect of *Ulva* extract is not apparent in an ABA-insensitive mutant

We next sought to determine whether ABA signalling could mediate the effects of *Ulva* extract on *Arabidopsis* development to uncover the mechanism by which *Ulva* extract inhibits germination. *Arabidopsis* seeds from the ABA-insensitive mutant *abi1* ^69,70^ were assayed for their response to *Ulva* extract. *abi1* seeds are unresponsive to the inhibitory effect of *Ulva* extract and behave similarly to untreated controls under all treatments (Fig. 2a-c). This suggests that the inhibition of *Arabidopsis* seed germination by *Ulva* extract depends on a functional ABA signalling pathway in the seeds.

**Figure 2.**
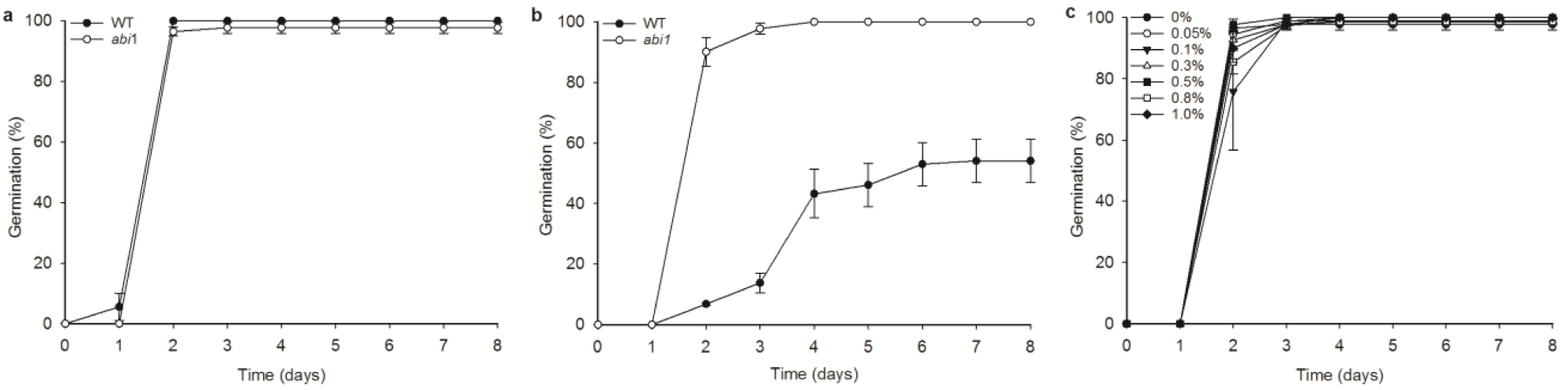
The *abi1* mutant is insensitive to the inhibitory effects of *Ulva* extract on germination. (a) Germination of WT and *abi1* on control medium. There are no significant differences between genotypes on any day (Kruskal-Wallis test). (b) Germination of WT and *abi1* on 1% *Ulva* extract. On days 2-8, wild-type is significantly (p<0.05, Kruskal-Wallis test) different from *abi1*. p=0.001 day2, p=0.001 day3, p=0.006 day4, p=0.005 day5, p=0.002 day6, p=0.001 day7, p=0.001 day8. (c) Germination of *abi1* seeds on increasing concentrations of *Ulva* extract. There are no significant differences between treatments apart from on day 2, when 0.5% and 0.8% treatments are significantly different from one another (p=0.023). The wild type data is the same as in Figure 1a. Bars represent standard error of the mean.

### The *abi1* mutant’s root growth responds normally to low concentrations of *Ulva* extract and is slightly insensitive to higher concentrations of *Ulva* extract

Since the *abi1* mutant is impaired in its germination response to *Ulva* extract and since ABA is known to have a biphasic effect on root growth ^65^, we tested the effect of *Ulva* extract on the root growth of the *abi1* mutant. The *abi1* mutant behaved similarly wild type plants at low concentrations (<0.1%) of *Ulva* extract (Figure 3 a, b). This suggests that the stimulatory effect of *Ulva* extract on root growth cannot be attributed to changes in ABA signalling in the plant. At higher concentrations (0.3% - 1%) of *Ulva* extract, the *abi1* mutant showed some insensitivity to inhibition of root growth (Figure 3c), but this was of a much smaller magnitude than the *abi1* mutant’s insensitivity during germination. This suggests that changes in ABA signalling in *Arabidopsis* may partially contribute to the inhibitory effect of *Ulva* extract on root growth.

**Figure 3.**
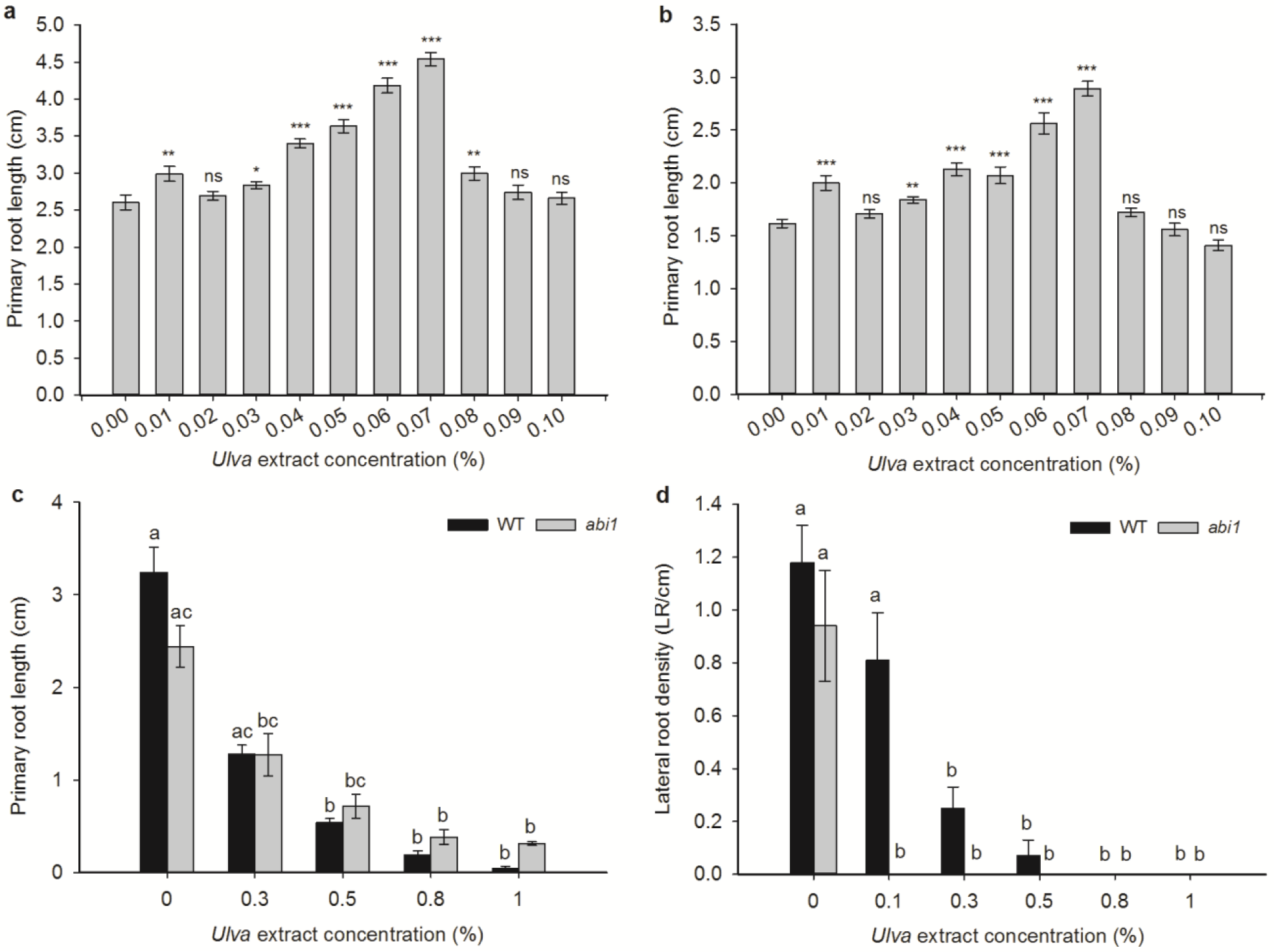
The *abi1* mutant is partially insensitive to the inhibitory effect of *Ulva* extract on root growth, responds to the stimulatory effect of *Ulva* extract on root growth similarly to wild-type, and is sensitive to the inhibition of lateral roots by *Ulva* extract. (a), (b) Comparison of wild-type (a) and *abi1* mutant (b) root-length responses to low concentrations (0-0.1%) of *Ulva* extract, measured on day 10. In wild-type, significant (p<0.05; Mann-Whitney U-test) differences are seen at 0.01% (p=0.0018), 0.03% (p=0.025), 0.04% (p=0.0000), 0.05% (0.0000), 0.06% (0.0000), 0.07% (p=0.0000, 0.08% (p=0.0074). In *abi1* mutants, significant differences are seen at 0.01% (p=0.0000), 0.03% (p=0.0003), 0.04% (p=0.0000), 0.05% (p=0.0001), 0.06% (p=0.0000), 0.07% (p=0.0000), 0.1% (p=0.0088). Asterisks: significant differences between treatments and 0% control: * p<0.05, ** p<0.01 and *** p<0.001. n=26-31 seedlings for each treatment and each genotype. (c) Comparison of wild-type and *abi1* mutant root-length responses to 0-1% *Ulva* extract, measured on day 10. Wild-type on 0% *Ulva* extract is significantly different to wild-type on 0.5%, 0.8% and 1% (all p<0.001). *abi1* on 0% *Ulva* extract is significantly different to *abi1* on 0.8 and 1% *Ulva* extract (p<0.001%). Wild-type on 0% is significantly different to *abi1* on 0.3% (p=0.010) and *abi1* on 0.5-1% (P<0.001%). *Ulva* extract treatments decrease root length in both wild-type and *abi1*. Between treatments, fewer significant differences are seen with *abi1* mutants than with wild-type plants, demonstrating *abi1*’s insensitivity to *Ulva* extract during root length inhibition (Kruskal-Wallis). Letters represent significant differences. n=15-20 seedlings for each treatment and each genotype. (d) Comparison of wild-type and *abi1* mutant lateral root responses to 0-1% *Ulva* extract calculated on day 10. Wild-type and *abi1* mutants are both inhibited by *Ulva* extract, with wild-type being significantly inhibited by 0.3% (p=0.001) and by 0.5%-1% *Ulva* extract (p<0.001) and *abi1* being significantly inhibited (p<0.001) by 0.1%-1% *Ulva* extract. Thus, *Ulva* extract treatments significantly decrease lateral root density in both wild-type and *abi1* (Kruskal-Wallis). n=30-60 seedlings for each treatment and each genotype; letters represent significant differences between genotypes and treatments. In all panels, bars represent standard error of the mean.

### The *abi1* mutant is sensitive to lateral root-inhibition by *Ulva* extract

Since ABA plays an inhibitory role in LR development ^71^, we next tested the effect of 0.1-1% *Ulva* extract on lateral root development in the *abi1* mutant. The *abi1* mutant’s LR development was inhibited by *Ulva* extract more strongly than wild type controls (Figure 3d), including at 0.1% *Ulva* extract, which has no effect on primary root growth. This implies that the inhibition of LR development by *Ulva* extract is not mediated by the ABA signalling pathway and that LRs respond differently to *Ulva* extract compared to the primary root.

### Elemental analysis of *Ulva intestinalis*

Although *Ulva* ^72^ and other seaweeds ^73^ are known to produce ABA, the *Ulva* extract used in our experiments is a water-soluble extract, so its effects on the *Arabidopsis* ABA signalling pathway are most likely indirect: ABA is soluble in organic solvents rather than water ^74,75^. We therefore measured the concentration of a panel of 31 water-soluble ions in our *Ulva intestinalis* samples using ICP-MS to determine whether (i) there were substances present in the tissue that were at markedly different levels to those in a land plant standard, (ii) whether any substances were present at substantially different levels to those in our standard *Arabidopsis* growth medium and (iii) whether the presence of any of the substances could explain the effects of *Ulva* extract on *Arabidopsis* seedling development. Our analysis identified 16 elements present at higher levels in *Ulva intestinalis* extract than in the land plant control, namely Boron, Sodium, Sulphur, Lithium, Aluminium, Vanadium, Manganese, Iron, Copper, Arsenic, Strontium, Silver, Caesium, Thallium, Lead and Uranium (Table 1). Of these, 12 elements (Sodium, Lithium, Aluminium, Vanadium, Copper, Arsenic, Strontium, Silver, Caesium, Thallium, Lead and Uranium) are present in 1% *Ulva intestinalis* extract at higher values than in *Arabidopsis* growth medium (Table 1). Out of these 12 elements, only three, namely Sodium, Aluminium and Copper, are present at levels known to have an effect on *Arabidopsis* development. The remaining 9 elements are present at micromolar (Lithium) or nanomolar quantities, while the published literature demonstrates their effects on *Arabidopsis* germination and root growth only in the micromolar to millimolar ^76-83^ range.

**Table 1.** Elemental analysis of *Ulva* intestinalis compared to land plant (tomato) control. Elements that show a significantly higher concentration in *Ulva* compared to tomato are highlighted in light grey. The concentration of each element in 0.05% *Ulva* extract (stimulates root growth) and 1% *Ulva* extract (inhibitory to germination and root growth) is shown, compared to the concentration of the same element in our normal *Arabidopsis* growth medium (0.5x MS). Elements highlighted in dark grey are present at higher concentrations in 1% *Ulva* extract than in 0.5MS.

| Element | Seaweed composition (mg/kg) | Tomato control composition (mg/kg) | Probability of same concentration in seaweed and tomato (Welch's t-test) | Concentration of element in 0.05% *Ulva* extract - stimulatory for root growth | Concentration of element in 1% *Ulva* extract - inhibitory for germination and root growth | Concentration of element in *Arabidopsis* medium |
| --- | --- | --- | --- | --- | --- | --- |
| B | **131.51±5.65** | 31.18±0.25 | 0.0046 | 6µM | 121µM | 100µM |
| Na | **61860.69±259.87** | 116.79±4.96 | 0.0258 | 530µM | **10.5mM** | **100µM** |
| Mg | 14291.19±1182.33 | 10178.93±71.28 | 0.1042 | 295µM | 5.9mM | 1.5mM |
| P | 2511.15±76.31 | 2307.47±15.21 | 0.1569 | 40µM | 800µM | 1.25mM |
| S | **24810.4±872.48** | 9841.569±110.91 | 0.0044 | 385µM | 7.7mM | >1.5mM |
| K | 16466.06±2383.77 | 27310.14±197.01 | 0.0642 | 210µM | 4.2mM | >19mM |
| Ca | 37045.93±3114.25 | 46397.44±256.12 | 0.1325 | 460µM | 9.2mM | 3mM |
| Ti | 18.13±0.36 | 18.79±0.13 | 0.2782 | 190nM | 3.8µM | N/A |
| Li | **2.88±0.19** | 0.5±0.02 | 0.0097 | 150nM | **3µM** | N/A |
| Be | 0.05±0.00 | 0.01±0.01 | 0.0967 | 3nM | 60nM | N/A |
| Al | **1397.88±177.11** | 478±2.71 | 0.0513 | 25.9µM | **518µM** | **N/A** |
| V | **2.85±0.22** | 0.74±0.00 | 0.0164 | 28nM | **560nM** | **N/A** |
| Cr | 2.38±0.23 | 1.29±0.24 | 0.1141 | 23nM | 460nM | N/A |
| Mn | **22.9±1.23** | 228.11±1.6 | 0.0003 | 210nM | 4.2µM | 100µM |
| Fe | **795.33±95.13** | 322.87±3.18 | 0.0056 | 5µM | 100µM | 100µM |
| Co | 0.34±0.04 | 0.47±0.00 | 0.1267 | 3nM | 60nM | 100nM |
| Ni | 1.68±0.13 | 1.4±0.1 | 0.2871 | 154.5nM | 290nM | N/A |
| Cu | **13.37±2.22** | 1.71±0.05 | 0.0504 | 105nM | **2.1µM** | **100nM** |
| Zn | 26.03±0.87 | 25.77±0.18 | 0.8336 | 200nM | 4µM | 30µM |
| As | **3.07±0.21** | 0.13±0.00 | 0.0080 | 20.5nM | **410nM** | **N/A** |
| Se | 0.13±0.01 | 0.07±0.00 | 0.1006 | 1nM | 20nM | N/A |
| Rb | 5.87±0.76 | 14.09±0.09 | 0.0117 | 34.5nM | 690nM | N/A |
| Sr | **125.03±1.47** | 86.5±0.29 | 0.0015 | 715nM | 14.3 µM | N/A |
| Mo | 0.34±0.04 | 0.44±0.02 | 0.156 | 2nM | 40nM | 1µM |
| Ag | **0.09±0.00** | 0.00±0.00 | 0.0001 | 415pM | **8.3 nM** | **N/A** |
| Cd | 0.19±0.01 | 1.48±0.00 | p<0.0001 | 1nM | 20nM | N/A |
| Cs | **0.28±0.033** | 0.05±0.00 | 0.0214 | 1nM | **20nM** | **N/A** |
| Ba | 10.37±0.91 | 64.04±0.24 | 0.0002 | 38nM | 760nM | N/A |
| Tl | **0.09±0.01** | 0.04±0.00 | 0.0278 | 220pM | **4.4nM** | **N/A** |
| Pb | **1.79±0.12** | 0.54±0.00 | 0.0138 | 4.5nM | **90nM** | **N/A** |
| U | **0.08±0.01** | 0.03±0.00 | 0.0327 | 170pM | **3.4nM** | **N/A** |

The level of Sodium in the 1% *Ulva* extract is 10.5mM, but *Arabidopsis* germination is inhibited only by concentrations of salt above 150mM ^84^. Thus, the level of germination-inhibition with *Ulva* extract at ≥0.3% is not attributable to the levels of Sodium in the extract. *Arabidopsis* root growth is inhibited by concentrations of 25mM Na^+^ and above ^85^. Thus, the inhibition of root growth seen in our experiments is unlikely to be attributable wholly to salt stress. This conclusion is in accordance with the fact that the *abi1* mutant is not wholly insensitive to the root growth inhibition (Fig. 3c) since salt stress responses are mediated by ABA signalling ^86,87^.

Aluminium (Al^3+^) ions are present in 1% *Ulva* extract at >500µM, and 0.05% *Ulva* extract at 26µM. Previous literature shows that even 5µM Al^3+^ can slow root growth ^88^ while 500µM Al^3+^ can reduce root growth by around 80% ^89^. Thus, the elevated Al^3+^ levels in the *Ulva* extract could be contributing to the inhibition in root growth that we see at concentrations of *Ulva* extract ≥0.3%. Copper ions (Cu^2+^) are present in 1% *Ulva* extract at 2.1µM (Table 1). Copper levels of 1.6µM have previously been described as rhizotoxic ^76^ and 20-25µM Cu^2+^ inhibits root elongation in several studies ^90^. Higher levels of copper (500µM-2mM) inhibit seed germination ^91,92^. Thus, it is unlikely that the elevated copper levels in the *Ulva* extract are causing germination inhibition, but they could be partly contributing to the inhibition in root growth that we see in ≥0.3% *Ulva* extract.

### Auxin, ethylene, cytokinin mutants respond similarly to wild-type *Arabidopsis* on *Ulva* extract with respect to root growth

Aluminium stress on roots is mediated by a combination of ethylene (via changes in auxin transport, at higher Al^3+^ concentrations) and cytokinin signalling (at lower Al^3+^ concentrations) ^89^. We tested mutants in auxin-, cytokinin and ethylene signalling for their root growth responses to *Ulva* extract (Supplemental Figure 1). The two auxin signalling mutants used were the receptor mutant *tir1-1* ^93^ and the auxin-resistant signalling mutant *axr1-1* ^94^. The two ethylene signalling mutants used were the receptor mutant *etr1-3* and the signalling mutant *ein2-1* ^95^. The cytokinin mutant used was the receptor mutant *cre1-1* ^45^. All mutants responded similarly to wild type seedlings to “inhibitory” concentrations of *Ulva* extract ranging from 0.1-1%, equating to approximately 50-500µM Al^3+^ (Supplemental Fig. 1a). This suggests that the ethylene, auxin and cytokinin hormone signalling pathways do not participate substantially in root growth inhibition by *Ulva* extract. Moreover, none of the mutants were insensitive to the root growth-stimulatory effect of *Ulva* extract (Supplemental Fig. 1b), suggesting that these hormones do not participate in the root growth stimulation brought about by *Ulva* extract. Similar data was obtained when the quintuple *della* mutant ^96^ was assayed for root growth stimulation and inhibition: the mutant behaved as wild-type, showing that gibberellin-DELLA signalling, which is involved in multiple plant stress- and growth-responses ^97^ is not involved in the effects of *Ulva* extract on root growth (Supplemental Figure 2).

### Cytokinin- and ethylene ethylene-signalling mutants show some insensitivity to inhibition of germination by *Ulva* extract

As Aluminium (Al^3+^) ions are present in 1% *Ulva* extract at >500µM, a concentration that causes very substantial decreases in root growth ^89^, and since two hormones involved in root responses to Aluminium, cytokinin and ethylene, are also regulators of seed germination ^98,99^, we also tested the germination behaviour of cytokinin receptor mutant *cre1* and the ethylene receptor mutant *etr1* on 0.1-1% *Ulva* extract. Both mutants’ seeds showed some insensitivity to germination-inhibition compared to wild type (Fig. 4a, 4b), but were not as insensitive as the *abi1* mutant (Fig. 2b). Both *cre1* and *etr1* also showed a higher final germination percentage in comparison to WT germination on 0.8% and 1% *Ulva* extract over the same period of time (Fig 1c, d, e). This suggests that the inhibition of *Arabidopsis* seed germination by *Ulva* extract is influenced by the cytokinin- and ethylene signalling pathways in addition to the ABA signalling pathway.

**Figure 4.**
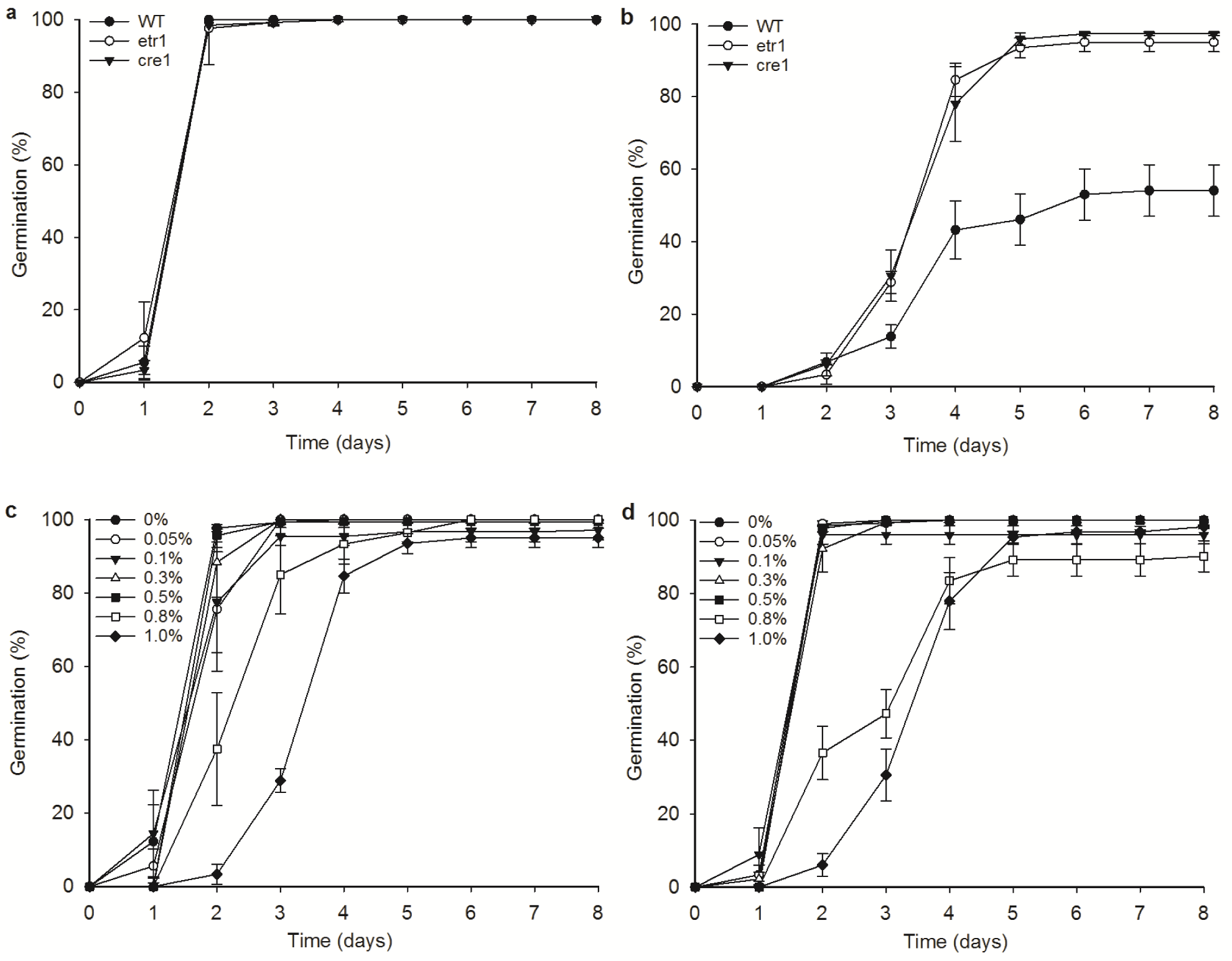
Ethylene and cytokinin signalling mutants are slightly insensitive to the inhibition of germination by *Ulva* extract. a) Wild type seed, *etr1* mutant seed and *cre1* mutant seed all germinate similarly on standard growth medium. No significant differences between genotypes are seen on any day. b) *etr1* and *cre1* mutant seed germinate faster than wild type in the presence of 1% *Ulva* extract. Significant (p<0.05%; Kruskal-Wallis) differences are seen between wild-type and *cre1* on day 5 (p=0.007) and on days 6-8 (each p=0.02). Significant differences between wild-type and *etr1* are soon on days 4 (p=0.028), 5 (p=0.009), 6 (p=0.004), 7 and 8 (each p=0.002). c) Germination of *cre1* mutant seed on varying concentrations of *Ulva* extract. On day 2, there is a significant (p<0.05%; Kruskal-Wallis) difference between 0.8% and all other treatments (p=0.035) and significant differences between 1% and control (p=0.015), 0.05% (p=0.007), 0.1% (p=0.025), 0.3% (p=0.015) and 0.5% (p=0.041). d) Germination of *etr1* mutant seed on varying concentrations of *Ulva* extract. On day 2, 1% *Ulva* extract is significantly different (p<0.05; Kruskal-Wallis) from the control (p=0.041), 0.05% (p=0.010), 0.1% (p=0.048), 0.3% (p=0.010) and 0.5% (p=0.035). On day 3, 1% Ulva extract is significantly different from control, 0.05% and 0.3% (all p=0.025). Wild type data in (a) and (b) is the same as in Figure 2.

## Discussion

*Ulva* extract at concentrations of 0.5%-1% inhibits wild-type *Arabidopsis* seed germination. *Ulva* extract ≥0.3% reduces wild-type *Arabidopsis* primary root growth and the extract inhibits wild type LR formation even at concentrations below 0.1%, suggesting that LRs are more sensitive than the primary root to the inhibitory agent(s) in the *Ulva* extract.

Our results concur with other studies where seaweed extract at high concentrations inhibited seed germination and seedling growth. Reduced germination occurred in pepper seeds primed with brown seaweed (*Ascophyllum*) extract at 1:250 (0.4%) and at higher concentrations (10%) of Maxicrop (commercial seaweed extract) solution compared to control seeds ^100^. A higher concentration (1.0%) of water-extracts from the brown seaweeds *Caulerpa sertularioides, Padina gymnospora* and *Sargassum liebmannii* reduced tomato germination and seedling development ^34^. 2%-10% aqueous extracts from *Sargassum johnstonii* led to similar effects on tomato ^101^.

Concentrations of kelp waste extracts (KWE) from 10–100% inhibited germination of pakchoi (*Brassica chinensis* L.). This was attributed to high levels of NaCl ^102^, which are absent from our *Ulva* extract. Arnon and Johnson (1940) reported detrimental effects on early tomato development as a result of higher pH in the growth medium. In our experiments, the pH was adjusted to be the same for all concentrations of *Ulva* extract so the effects are not due to altered pH.

*Ulva* extract has a growth-stimulating effect on wild type *Arabidopsis* primary root elongation specifically at concentrations between 0.025-0.08%. This is in accordance with data from other species, suggesting that *Arabidopsis* is a good model for studying the effects of seaweed extracts. Seaweed extract may improve water and nutrient uptake efficiency by root systems ^103^ leading to enhanced plant growth and vigour. Commercial extracts from the brown seaweed *Ecklonia maxima* stimulated tomato root growth at low concentrations (1:600; 0.17%) while higher concentrations (1:100; 1%) strongly inhibited root growth ^33^. Root growth enhancement was seen in *Arabidopsis* plants treated with aquaeous *Ascophyllum nodosum* extracts (0.1gL^-1^; 0.01%), whereas plant height and number of leaves were affected positively at 1gL^-1^ (0.1%) ^104^. Lower concentrations (0.2%) of extracts of both *Ulva lactuca* (green seaweed) and *P. gymnospora* (brown) were more effective at enhancing tomato seed germination (Hernández-Herrera et al., 2014). We observed no boost in *Arabidopsis* germination with *Ulva intestinalis* extract under our growth conditions where we vernalise seeds to break dormancy before an assay, so this may explain the discrepancy between the experiments.

2% kelp waste extract (KWE) stimulated pakchoi seed germination ^102^. Pakchoi seedling growth (plumule length, radicle length, fresh weight and dry weight) was improved by treatment with 2-5% KWE ^102^. This data is in-line with our observed root growth stimulation at low concentrations. The KWE was prepared differently (cell wall digestion and centrifugation) to our *Ulva intestinalis* extract, which may explain why higher concentrations of KWE than *Ulva* extract give stimulatory effects; in addition, pakchoi is larger than *Arabidopsis*.

The stimulatory effect of KWE on pakchoi may be attributed to the combined effects of soluble sugars, amino acids and mineral elements ^102^. Sugars are immediate substrates for intermediary metabolism and effective signaling molecules: thus accessibility of sugars influences plant growth and development ^105^. The growth-enhancing potential of algal extract correlates with the presence of diverse polysaccharides, including unusual/complex polysaccharides not present in land plants ^21,106^. However, a role for macro- and microelements, vitamins and phytohormones is also suggested ^20,27,32,107-109^.

Since our *Ulva* extracts are water-based, it is unlikely that they contain high quantities of plant hormones, which are largely soluble in organic solvents. Our *Arabidopsis* mutant analysis demonstrates that germination-inhibition by *Ulva* extract is dependent on activation of the *Arabidopsis* ABA signaling pathway, with cytokinin- and ethylene-signaling also playing a role. This suggests that a substance(s) in *Ulva* extract activates endogenous plant hormone signaling to inhibit germination. *Ulva* extract-mediated inhibition of primary root growth is partly blocked in an ABA-insensitive mutant, while cytokinin-, auxin-ethylene- and gibberellin signaling mutants all respond similarly to wild type with respect to root growth. This implies that although ABA signaling plays a role in primary root growth inhibition by *Ulva* extract, additional pathways also contribute. Lateral root development is inhibited via a different mechanism to primary root growth, as the ABA-insensitive *abi1* mutant’s LR development is inhibited by *Ulva* extract to a greater extent than wild-type (Fig. 3d).

Our elemental analysis of *Ulva* tissue suggests that the most likely cation contributing to the inhibitory effects of *Ulva* extract is Al^3+^, which is present in quantities known to inhibit *Arabidopsis* primary root growth ^88,89^. Al^3+^ may not be the only inhibitory substance present: previous research has demonstrated a role for auxin, ethylene and cytokinin in root responses to Al^3+^ stress ^89^ and this is not apparent from our mutant root assays. Conversely, there may be other hormones involved in seed- and root responses to Al^3+^ stress: the effects of Al^3+^ on germination and lateral root development in *Arabidopsis* has not previously been studied. The toxic effect of Al^3+^ in the *Ulva* extract may be partially countered by the relatively high levels of Mg^2+^ also present in the extract (In 1% *Ulva* extract, 4x that present in *Arabidopsis* growth medium - Table 1; ^79^).

Al^3+^ stress has a range of physiological effects that could affect root growth and development. Al^3+^ stress alters membrane potentials, which affects transport of ions, including Ca^2+^, across membranes. This can result in changes in cytoplasmic Ca^2+^ homeostasis, which controls cell signaling, metabolism and cell-growth processes including root development ^110^. Al^3+^ stress induces changes in the expression and activity of the plasma membrane H^+^-ATPase that controls cytosolic pH and membrane potentials ^111^.

Seaweeds contain high levels of particular cations: macroelements (Na, P, K, Ca) and microelements (Fe, B, Mn, Ca, Mo, Zn, Co) that have critical roles in plant development and growth ^112,113^. In many vegetable crops, the accumulation of sodium ions restrains embryo or seedling development, leading to reduced germination, uneven morphogenesis and loss of crop production e.g. ^114^. Our data suggests that the only macroelement present at higher concentrations in *Ulva* extract than in plant tissues (or indeed plant growth medium) is Na^+^, but Na^+^ is not present at high enough concentrations to explain the inhibition of germination, root growth and lateral root development that we see. *Ulva* species tolerate low salinity despite being marine algae. Our *Ulva* sampling site is where a river meets the sea: the salinity of the seawater is low (F. Ghaderiardakani, unpublished). A reduction in germination rate and growth of tomato attributable to salt (and perhaps reduced imbibition of water by seeds) was suggested upon applying brown seaweed (*Caulerpa sertularioides* and *Sargassum liebmannii*) liquid extracts, but not with *U. lactuca* and *P. gymnospora* with a lower salt concentration ^34^.

Some seaweed extracts alleviate salt stress: the survival of Kentucky bluegrass (*Poa pratensis* L. cv. Plush) treated with a proprietary seaweed extract (38Lha^-1^) increased significantly, under various levels of salinity, with improved growth and promotion of rooting of the grass at a soil salinity of 0.15Sm^-1^ ^19^. Application of seaweed extract activated a mechanism reducing the accumulation of Na^+^ in plants; grass treated with seaweed extract had less sodium in the shoot tissue ^115,116^.

The microelements B and Fe are present at higher concentrations in *Ulva* tissue than in our land plant control, but at levels that are very similar to that found in our *Arabidopsis* growth medium, so cannot be contributing to the observed stimulatory or inhibitory effects. The content of minerals in *Ulva intestinalis* is in-line with values for *Ulva* spp. reported previously, e.g. *Ulva lactuca* ^34^ and *Ulva reticulata* ^38,112^.

Using seaweed extracts as biofertilisers due to their direct or indirect stimulatory impacts on plant metabolism has been suggested as one of their key beneficial applications ^23^. Taken together, our results and others’ suggest that for plants to benefit optimally from algal extracts, only a small quantity should be used or could be mixed with commercially available fertilisers for a synergistic effect on crop yield and a reduction in quantities and costs of chemical fertilisers applied ^117^.

Our data demonstrates that *Ulva* extract can inhibit *Arabidopsis* seed germination, early root growth and lateral root development, even at concentrations below 1%, by activating endogenous plant hormone signaling pathways. Could this in itself be useful? One of the top priorities in organic agriculture is the eradication of weeds from the production area ^118^. Concerns about improvements in agriculture focus on diminishing weeds’ adverse effects on the environment and improving the sustainable development of agricultural systems. New approaches are required to integrate biological and ecological processes into food production and minimize the use of practices that lead to the environmental harm ^119^. Considering the observed biological inhibitory effects resulting from the action of seaweed extracts on crops’ germination and early development particularly at high concentration, it might be worthwhile to employ seaweed extracts as organic herbicides. The evidence at hand establishes that there are benefits to be obtained from utilizing macroalgal products in agricultural systems. Further translational studies are required to define the appropriate algal sources for commercial biostimulants (considering inherently different algal extracts and also the availability of seaweed biomass in a particular area), their application form and frequency, the timing of applications in relation to plant development and the optimal dosages needed to maximise both agricultural productivity and economic advantages.

In conclusion, water-soluble algal extracts from *Ulva intestinalis* were effective at stimulating the primary root growth of *Arabidopsis thaliana* only when applied at low concentrations. High concentrations of *Ulva* extract inhibit germination and root development, perhaps in part due to Al^3+^ toxicity, with endogenous plant ABA signalling playing a role in this inhibition. The effects of algal extracts on *Arabidopsis* development are likely mediated by a complex interplay of hormones. Future work targeting candidate genes in *Ulva* ^60^ may uncover how *Ulva* extracts exerts their effects on plant hormone signalling. Although if used sparingly, seaweed extracts are potential candidates to produce effective biostimulants, they may be just as beneficial as organic herbicides by targeting plants’ ABA signalling mechanisms. Cross-disciplinary research could help farmers to benefit optimally from the use of algal extracts in the future, particularly for cost-effective organic farming and an environmentally-friendly approach for sustainable agriculture.

## Methods

### Collection and Identification of Algal Samples

Vegetative and fertile *U. intestinalis* blades were collected from the intertidal zone at low tide, three times between March 2015 and April 2016, from the coastal area of Llantwit Major beach, South Wales, UK (51°40’ N; 3°48’ W). Excess water and epiphytic species were removed at the site by blotting the sample’s surface before storage on ice for transport back to the laboratory. Epiphyte-free samples were subjected to a molecular identification using plastid-encoded *rbcL* (large unit ribulose bisphosphate carboxylase) and *tuf*A (plastid elongation factor) markers as identification solely by morphological characteristics is not reliable ^61^.

### Preparation of water-soluble *Ulva* Extract

*Ulva* samples were washed with tap water to remove surface salt, shade dried for 10 days, oven-dried for 48h at 60 °C, then ground to a fine powder using a coffee grinder (Crofton, China) to less than 0.50mm. 10g of this milled material was added to 100mL of distilled water with constant stirring for 15 min followed heating for 45 minutes at 60°C in water bath ^38^. The contents were filtered through two layers of muslin cloth. This *Ulva* extract was designated as 10% stock solution and added to MS solution to make up specific concentrations and autoclaved. 1% *Ulva* extract stock was subjected to pH measurement and elemental analysis. All measurements were performed in triplicate.

### Digestion of plant material for elemental analysis

*Ulva* samples were digested using a microwave system, comprising a Multiwave 3000 platform with a 48-vessel MF50 rotor (Anton Paar GmbH, Graz, Austria); digestion vessels comprised perfluoroalkoxy liner material and polyethylethylketone pressure jackets (Anton Paar GmbH). Dried samples (∼0.2g) were digested in 2mL 70% Trace Analysis Grade HNO3, 1 mL Milli-Q water (18.2 MΩ cm; Fisher Scientific UK Ltd, Loughborough, UK), and 1mL H2O2 with microwave settings as follows: power =1400 W, temp = 140 C, pressure = 20 Bar, time = 45 minutes. Two operational blanks and two certified reference material of leaf (Tomato SRM 1573a, NIST, Gaithersburg, MD, USA) were included in each digestion run. Following digestion, each tube was made up to a final volume of 15mL by adding 11mL of Milli-Q water and transferred to a universal tube and stored at room temperature.

### Elemental analysis

Sample digestates were diluted 1-in-10 using Milli-Q water prior to elemental analysis. The concentrations of 28 elements were obtained using inductively coupled plasma-mass spectrometry (ICP-MS; Thermo Fisher Scientific iCAPQ, Thermo Fisher Scientific, Bremen, Germany); Ag, Al, As, B, Ba, Ca, Cd, Cr, Co, Cs, Cu, Fe, K, Mg, Mn, Mo, Na, Ni, P, Pb, Rb, S, Se, Sr, Ti, U, V, Zn. Operational modes included: (i) a helium collision-cell (He-cell) with kinetic energy discrimination to remove polyatomic interferences, (ii) standard mode (STD) in which the collision cell was evacuated, and (iii) a hydrogen collision-cell (H_2_-cell). Samples were introduced from an autosampler incorporating an ASXpress™ rapid uptake module (Cetac ASX-520, Teledyne Technologies Inc., Omaha, NE, USA) through a PEEK nebulizer (Burgener Mira Mist, Mississauga, Burgener Research Inc., Canada). Internal standards were introduced to the sample stream on a separate line via the ASXpress unit and included Sc (20µgL^-1^), Rh (10µgL^-1^), Ge (10µgL^-1^) and Ir (5µgL^-1^) in 2% trace analysis grade HNO_3_ (Fisher Scientific UK Ltd). External multi-element calibration standards (Claritas-PPT grade CLMS-2; SPEX Certiprep Inc., Metuchen, NJ, USA) included Ag, Al, As, B, Ba, Cd, Ca, Co, Cr, Cs, Cu, Fe, K, Mg, Mn, Mo, Na, Ni, P, Pb, Rb, S, Se, Sr, Ti (semi-quant), U, V and Zn, in the range 0 – 100µgL^-1^ (0, 20, 40, 100µgL^-1^). A bespoke external multi-element calibration solution (PlasmaCAL, SCP Science, Courtaboeuf, France) was used to create Ca, K, Mg and Na standards in the range 0-30mgL^-1^. Boron, P and S calibration utilized in-house standard solutions (KH_2_PO_4_, K_2_SO_4_ and H_3_BO_3_). In-sample switching was used to measure B and P in STD mode, Se in H_2_-cell mode and all other elements in He-cell mode. Sample processing was undertaken using Qtegra™ software (Thermo Fisher Scientific) with external cross-calibration between pulse-counting and analogue detector modes when required ^120^. Differences between seaweed and tomato control were analysed using a Welch’s t-test.

### Germination Bioassay

*Arabidopsis thaliana* wild-type Col-0 and mutant lines *abi1-1, tir1-1, axr1-3, cre1-12, etr1-3*, and *ein3-1* were obtained from the Nottingham Arabidopsis Stock Centre (Loughborough, UK). *Arabidopsis* seeds were sterilised in 20% Parozone^™^ bleach on a turning wheel for 10 minutes and subsequently washed 2-3 times in sterile water. Seeds were vernalized at 4°C for 48h and placed on 1% agar, containing 0.5MS and *Ulva* extract. Plates were transferred to the growth room for 7-10 days and incubated at 22 ± 2°C with a 16-h-light photoperiod and light intensity of 120µmolm^-2^s^-1^. Germination was observed daily as in ^121^. A seed was scored as germinated when its radicle had emerged from within the seed coat. Germination percentage (GP) was calculated as follows: GP = (the number of germinated seeds/total number of seeds) ×100). Data from three independent biological repeats (n=30-90 seeds per genotype and treatment) were combined. To identify significant differences between treatments and genotypes, Kruskal-Wallis one-way ANOVA on ranks followed by Dunn’s post-hoc tests were performed using SigmaPlot 13 software (Systat Software, San Jose, CA).

### Root Bioassay

Experiments were conducted using 10cm square agar plates. 20 seeds were placed individually on the agar following a line across the top of the plate. The plates were sealed with Micropore tape (3M), taped together and incubated vertically in standard growth conditions as in ^121^.

From day 7 to 14 the seedlings were photographed and primary root (PR) lengths were measured with ImageJ open-source software (http://rsb.info.nih.gov/ij). For some assays, the number of visible emerged lateral roots (LR) on each primary root was also counted and the lateral root density was calculated by dividing the number of LRs present by the length of that root. To identify significant differences between treatments and controls in wild-type plants, data were first checked to confirm normality, then appropriate two-tailed t-tests (normal data) or Mann-Whitney U-tests (non-normal data) were performed in Excel using an Excel template from Gianmarco Alberti’s lab (xoomer.alice.it/Exceltemplates.pdf), comparing the results of each *Ulva* extract concentration to the control (without *Ulva* extract). To identify significant differences between treatments and genotypes, Kruskal-Wallis one-way ANOVA on ranks followed by a Dunn’s post-hoc test were performed using SigmaPlot 13 software (Systat Software, San Jose, CA). All experiments were repeated a minimum of two and a maximum of four times with similar trends observed in each biological repeat.

## Acknowledgements

This work was funded by an Islamic Development Bank PhD scholarship to FG and University of Birmingham funds for JC to host EC, DKD and KT in the lab.

## Author contributions

FG and JCC conceived the study and designed experiments. All authors performed experiments and analysed data: FG - Fig 1a,d,e, Fig 2, Fig 3a,b,c,d; Fig 4; EC - Fig 1a,c, Supplemental Fig 1; Fig 4; DKD – Fig 1e, Fig 3c,d, Supplemental Fig 1; KT – Fig 1a; Fig 2; NSG - Table 1, JCC – Supplemental Fig 2. FG supervised EC, DKD, KT in the lab; JCC supervised FG, ED, DKD, KT. FG, NSG and JCC wrote the paper.

## Data availability statement

The datasets generated during the current study are available from the corresponding author on reasonable request.

---

**Supplemental Figure 1.**
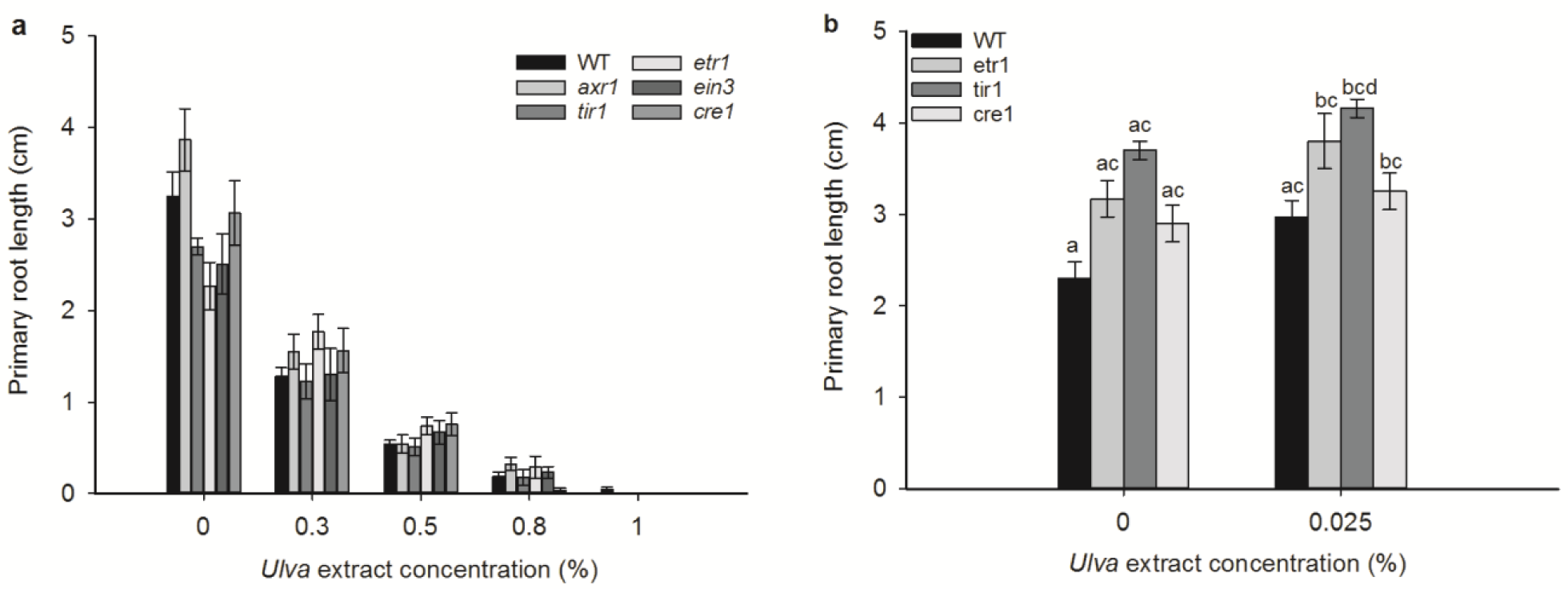
Ethylene, auxin and cytokinin signaling mutants’ root growth is inhibited by high concentrations of *Ulva* extract, similarly to wild-type plants. a) Inhibition of root growth by 0.3-1% *Ulva* extract in all hormone mutants, similarly to wild type. Comparison across genotypes and treatments using a Kruskal-Wallis test and a Dunn’s post-hoc test indicates that 0.3% *Ulva* extract reduces root growth (p≤0.05%) compared to the control in all genotypes but *etr1*, while 0.5% reduces root growth (p≤0.05%) compared to the control in all genotypes but *cre1*. 0.8% and 1% *Ulva* extract significantly reduce root growth in all genotypes compared to the control (p≤0.05). In terms of between-genotype differences, under control conditions only *etr1* is significantly different from other genotypes (p<0.05), at 0.3% and 0.5% there are no significant differences between the genotypes, at 0.8% *cre1* is significantly different from wild-type and *axr1* (p<0.05) and at 1% no genotypes are significantly different from one another. b) Stimulation of root growth in all hormone signaling mutants by 0.025% *Ulva* extract. An analysis of variance followed by a Tukey’s post-hoc test demonstrates significant differences between wild-type control and *etr1* and *tir1* treated with *Ulva* extract (p<0.01), between wild type control and cre1 treated with *Ulva* extract (p<0.05), between wild-type treated with *Ulva* extract and and *tir1* treated with *Ulva* extract (p<0.05), between *cre1* control plants and *etr1* treated with *Ulva* extract, and between *cre1* control and *tir1* treated with *Ulva* extract. These differences are represented by the letters above the graph.

**Supplemental Figure 2.**
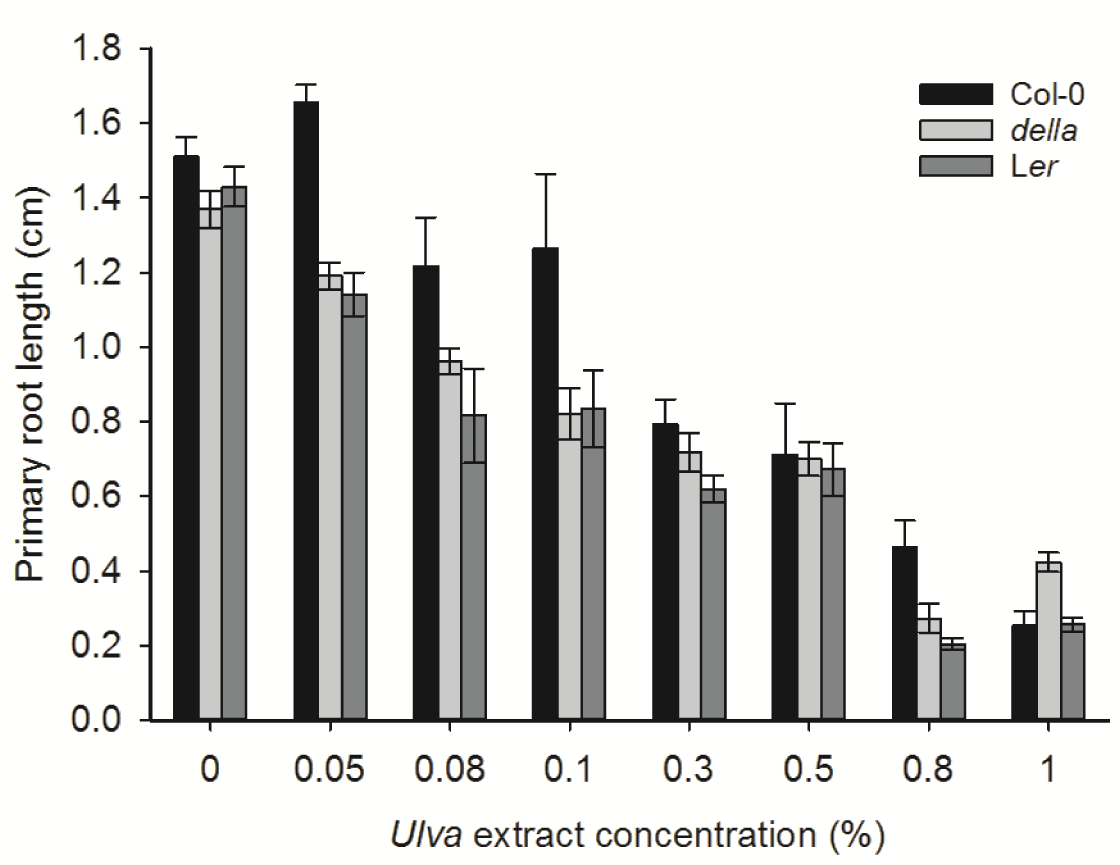
The *Arabidopsis* quintuple *della* mutant is sensitive to the inhibitory effects of *Ulva* extract on root growth. The *della* mutant was generated in a L*er* background also containing 3Mb of Col-0 sequence ^96^. For each *Ulva* extract concentration, the effects on Col-0, *della* and L*er* are shown.

